# Oxygen concentration modulates colibactin production

**DOI:** 10.1101/2022.06.20.496773

**Authors:** Nadège Bossuet, Cécile Guyonnet, Camille V Chagneau, Min Tang-Fichaux, Marie Penary, Dorian Loubet, Priscilla Branchu, Eric Oswald, Jean-Philippe Nougayrede

## Abstract

Up to 25% of the *E. coli* strains isolated from the feces of healthy humans harbor the *pks* genomic island encoding the synthesis of colibactin, a genotoxic metabolite. Evidence is accumulating for an etiologic role of colibactin in colorectal cancer. Little is known about the conditions of expression of colibactin in the gut. The intestine is characterized by a unique oxygenation profile, with a steep gradient between the physiological hypoxic epithelial surface and the anaerobic lumen, which favors the dominance of obligate anaerobes. Here, we report that colibactin production is maximal under anoxic conditions and decreases with increased oxygen concentration. We show that the aerobic respiration control (ArcA) positively regulates colibactin production and genotoxicity of *pks+ E. coli* in response to oxygen availability. Thus, colibactin synthesis is inhibited by oxygen, indicating that the *pks* biosynthetic pathway is adapted to the anoxic intestinal lumen and to the hypoxic infected or tumor tissue.

## Introduction

*E. coli* strains isolated from the feces of healthy subjects or from extra-intestinal infections, frequently harbor the *pks* genomic island that allow synthesis of a genotoxin called colibactin (1). This 54 kb *pks* island encodes a complex biosynthesis pathway, with polyketide synthases (PKS) and nonribosomal peptide synthetases (NRPS) as well as maturation enzymes, which cooperatively synthesize colibactin (2). Colibactin is an unstable peptide-polyketide metabolite, sensitive to aerobic oxidation and subsequent inactivation (3). Although colibactin is not directly purifiable, its bioactivity has been demonstrated in infected epithelial cells or on DNA exposed to *pks*+ bacteria: colibactin binds covalently to both strands of the DNA helix, resulting in interstrand crosslinks (4–6). Because these DNA lesions are highly toxic, the *pks* island also encodes self-protection systems (such as the ClbS protein) to protect the producing bacterium from the toxin (7–9). Colibactin is also toxic to the bacteria in the microbial community (10, 11) and to host mammalian cells. Epithelial cells exhibit DNA damage, cell cycle arrest, senescence and death upon exposure to *pks*+ bacteria (12, 13). Colibactin induces apoptotic cell death in lymphocytes and promotes virulence of *pks+ E. coli* during sepsis and meningitis (14–16). Colibactin is also produced in the gut by *pks*+ *E. coli* and inflicts DNA damage in intestinal epithelial cells (17–19). This DNA damage can ultimately lead to gene mutagenesis, and colibactin is suspected of promoting colon cancer (2, 17, 18). In mouse models that recapitulate intestinal tumorigenesis, inflammation is essential for cancer promotion by *pks+ E. coli*, and inflammation increases the transcription of *pks-*encoded genes (18, 20, 21). Thus, these bacteria appear to promote cancer under specific environmental conditions that favor expression of colibactin.

Recent studies have highlighted how environmental conditions regulate colibactin expression in *pks+ E. coli*. Indeed, it has been shown that colibactin production is regulated by iron availability (22). Factors originating from the producing bacterium or the host diet also modulate genotoxin production. The polyamine spermidine from bacterial metabolism or the environment is required for colibactin production (23). Similarly, carbon source impacts *pks* gene expression through regulation of central carbon metabolism (24). Oligosaccharides known to regulate bacterial metabolism increase colibactin expression, but this effect is abrogated by ferrous sulfate supplementation (25). Together, these studies suggest that colibactin production is constrained by the conditions in the bacterial microniche in the intestinal lumen.

The concentration of molecular oxygen (O_2_, hereafter referred to as oxygen) is a key environmental factor in the gut, playing a central role in metabolic processes of both the host and intestinal bacteria. The intestinal niche exhibits low oxygen tension, with a steep gradient from anoxia in the lumen to physiological hypoxia at the epithelial surface (26). Oxygen serves as an environmental cue for *E. coli*, with the Anoxic Redox Control (or Aerobic Respiration Control; Arc) system detecting oxygen deprivation to regulate the expression of anaerobic adaptation genes (27). The physiological hypoxia in the gut can be disrupted during disease, such as at sites of mucosal inflammatory lesions, or in tumors, which can be deeply hypoxic (28, 29). On the other hand, chronic inflammation results in increased epithelial oxygenation favoring aerobic proliferation of *E. coli* (30, 31). This has led to the hypothesis that oxygenation of the intestinal niche is an important factor in the carcinogenic activity of *pks+ E. coli* (31). The possibility that oxygen may directly modulate colibactin production by *pks+ E. coli* has not yet been studied. In this paper, we report the serendipitous observation that oxygenation of the bacterial culture has a marked impact on the toxicity of the genotoxin on the producing bacterium. We further demonstrate that oxygen inhibits colibactin production and genotoxicity in infected epithelial cells, and that the Arc system is a positive regulator of colibactin production.

## Results

### Oxygen inhibits colibactin autotoxicity in a colibactin-producing *E. coli clbS* mutant

The colibactin biosynthetic pathway encodes self-protective systems, including the ClbS protein that hydrolyzes colibactin and protects the *E. coli* genome DNA from colibactin toxicity (7, 8). We previously observed that a laboratory K12 strain of *E. coli* transformed with a BAC harboring the *pks* island with a deleted *clbS* gene exhibits an autotoxicity phenotype, with impaired growth compared to the control strain harboring the intact *pks* island (8). While studying this *clbS* mutant, we fortuitously observed that its autotoxicity phenotype was sensitive to specific culture conditions. In particular, we noticed that the use of culture tubes with tightly or loosely closed caps resulted in autotoxicity or normal growth, respectively. Based on this observation, we found that the *clbS* mutant showed a significant ∼10-fold decrease in CFU count compared with wild-type or complemented mutant strains when grown in small closed tubes, but not in large open tubes (Figure 1A). We thought that differences in culture oxygenation were a likely explanation for this observation. To confirm a role of oxygen in the expression of the *clbS* mutant autotoxicity phenotype, we tested cultures under atmospheric or hypoxic conditions. Culture tubes with vented caps were incubated with agitation in a standard atmosphere incubator (21% O_2_) or in a dedicated hypoxic chamber with a 0.1% O_2_ atmosphere. In the hypoxic chamber, the *clbS* mutant showed a marked decrease of approximately 100-fold in CFU numbers compared with the wild type or the complemented mutant (Figure 1B). In contrast, the *clbS* culture in the 21% O_2_ incubator did not show such a decrease in growth.

**Figure 1:**
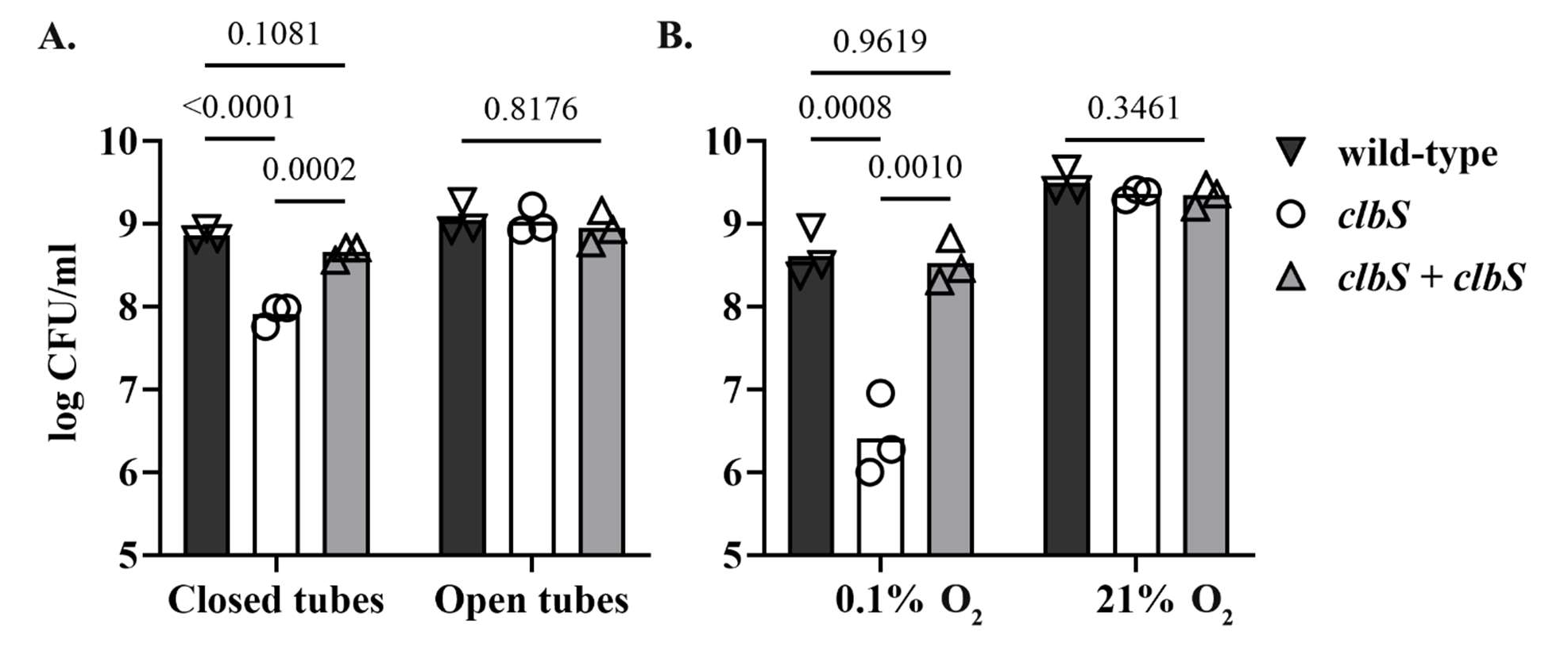
Culture oxygenation modulates the autotoxicity phenotype of a *clbS* mutant in a *pks*+ *E. coli*. **A**. *E. coli* strain DH10B pBAC*pks*, the isogenic *clbS* mutant deficient for colibactin self-resistance protein, and the trans-complemented *clbS* mutant (*clbS* + *clbS*), were grown (5 ml LB) with shaking in a standard atmospheric incubator, in 10 ml tubes that were tightly closed, or in 50 ml tubes with the cap opened. **B.** The bacteria were cultured with shaking in tubes with cap in vented position, within a hypoxic chamber at 0.1% O_2_, or in a standard atmosphere incubator (21% O_2_). After 16 h culture, bacterial growth was determined by plating and counting colony forming units (CFU). The mean and individual results of three independent experiments are shown, with the p values of an ANOVA and Tukey’s multiple comparison test.

### Oxygen inhibits DNA-crosslinking by *pks*+ *E. coli*

The reduced autotoxicity phenotype in the *clbS* mutant exposed to high oxygen concentration suggested that colibactin production, and/or its genotoxic activity, was inhibited. We therefore examined the effect of various oxygen concentrations on colibactin production and activity. Mature colibactin cannot be purified, but its genotoxicity can be detected by its DNA cross-linking activity on exogenous DNA exposed to *pks+ E. coli* bacteria (4). We used the human clinical *pks+ E. coli* strain SP15, which has been previously shown to produce colibactin in the mouse intestinal lumen (17, 31) and to alter the gut microbiota through the production of colibactin (32). DNA cross-linking activity of *E. coli* strain SP15 on linear DNA was tested at different oxygen concentrations (0-21%). DNA cross-linking by colibactin was readily detected at low (<1%) oxygen concentrations but decreased markedly at higher (4 and 13%) concentrations, becoming undetectable at 21% O_2_ (Figure 2AB). The growth of *E. coli* strain SP15 was similar regardless of oxygen concentration (Figure 2C), indicating that the modulation of its cross-linking activity was not explained by the number of bacteria. Thus, oxygen inhibits the activity and/or production of colibactin.

**Figure 2:**
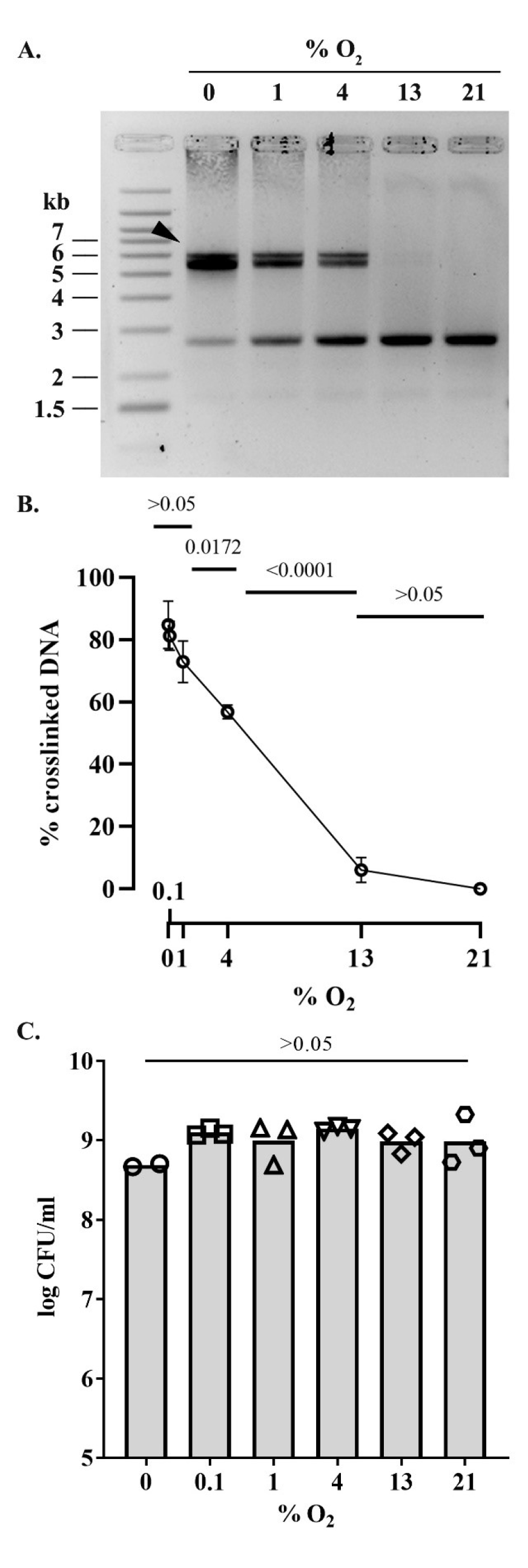
Effect of oxygen concentration on the DNA crosslinking activity of a *pks+ E. coli*. **A.** Linearized plasmid DNA was exposed 40 min to the *pks*+ *E. coli* strain SP15 grown 3.5 hours with shaking in vented cap tubes within an incubator regulated at the given percentage of oxygen, and then analyzed by denaturing gel electrophoresis. The sizes of the DNA fragments in the ladder are noted in kilobase. The DNA cross-linked by colibactin with apparent doubling in size compared to the intact denatured DNA is shown with an arrow. **B.** The percentage of DNA signal in the upper, crosslinked DNA relative to the total DNA signal in the lane was determined by image analysis. The mean and standard error of three independent experiments are shown, with the p values of an ANOVA and Tukey’s multiple comparison test. **C.** In the same experiments, the bacterial growth following 3.5 hours culture at given percentages of oxygen was examined by plating and counting colony forming units (CFU).

### Oxygen inhibits the expression and production of colibactin by *pks+ E. coli*

To examine whether oxygen modulates colibactin production, we quantified the metabolite N-myristoyl-D-AsnOH (C14-Asn), a stable byproduct released during the final step of colibactin synthesis (9). *E. coli* strain SP15 was grown as before with different oxygen concentrations, lipids were extracted from the culture supernatants and analyzed by quantitative chromatography coupled to mass spectrometry. C14-Asn production by SP15 was highest at low oxygen concentrations (<1% O_2_) and decreased at higher concentrations, with only background levels at 13 and 21% O_2_ (Figure 3). Thus, the synthesis of colibactin was the highest at low oxygen concentration but diminished with increasing oxygen concentration.

**Figure 3:**
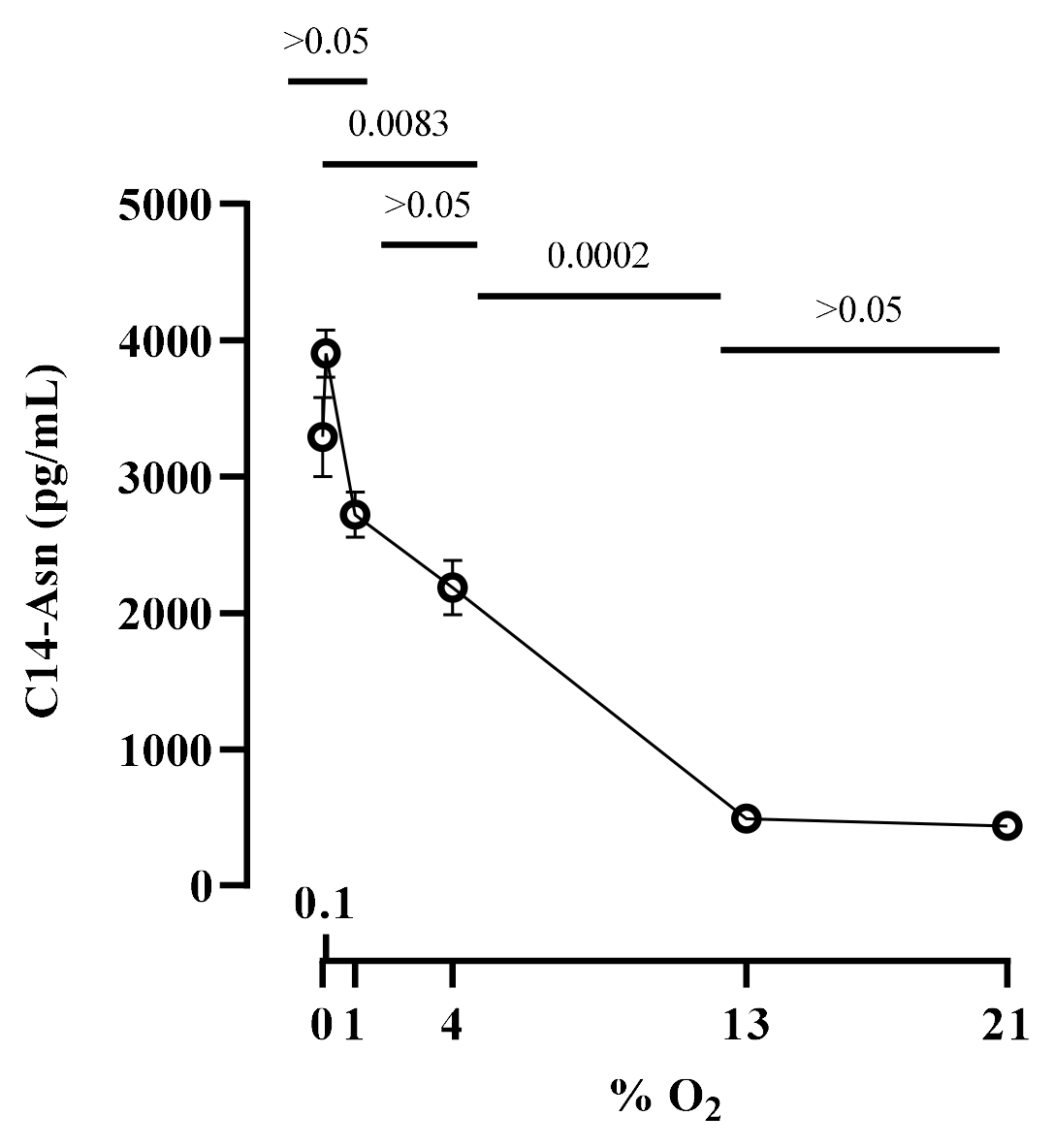
Effect of oxygen concentration on the production of colibactin cleavage product C14-Asn. The *E. coli* strain SP15 was grown 3.5 hours with shaking in vented tubes in an incubator regulated at various percentage of oxygen, then the culture supernatants were collected, the lipids were extracted and colibactin cleavage product C14-Asn was quantified by liquid chromatography coupled to mass spectrometry. The mean and standard error of three independent cultures are shown, with the p values of an ANOVA and Tukey’s multiple comparison test. The error bars of the biological triplicate samples at %O_2_>=13% are too small to appear on the graph.

To confirm that oxygen inhibits colibactin synthesis, we measured the activity of the promoter of *clbB*, which encodes one of the key NRPS-PKSs in the synthesis pathway (33). *E. coli* strain SP15 harboring a luciferase reporter system under the control of either a known constitutive promoter or the *clbB* promoter was grown with different oxygen concentrations, and then luminescence was quantified (Figure 4). The constitutive promoter showed no significant change in activity at all oxygen concentrations, whereas the *clbB* promoter showed its highest activity in anoxic cultures, and decreasing activity with increasing oxygen concentration (Figure 4). This result is consistent with a recent report of increased transcription of a *clbB-lacZ* reporter under oxygen-limited conditions (10). To confirm this finding, we measured by quantitative PCR the expression of the *clbB* gene and that of a control reference gene. We found that *clbB* mRNA expression was highest in anoxic cultures and decreased with increasing oxygen concentration (Figure 5). Together with the observation that DNA cross-linking activity and colibactin autotoxicity are inhibited by oxygen, these results demonstrated that oxygen constrains colibactin production.

**Figure 4:**
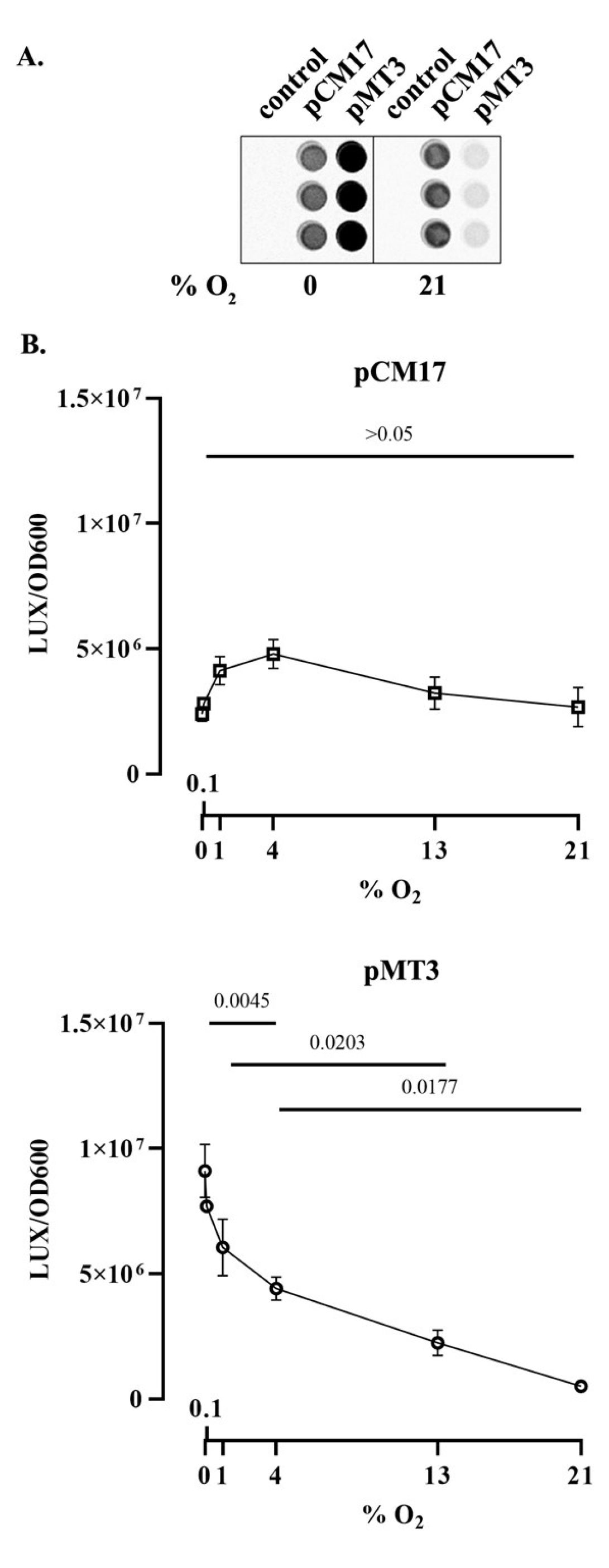
Effect of oxygen concentration on *clbB* promoter activity. **A.** Luminescence of *E. coli* strain SP15 carrying the luciferase reporter plasmid pCM17 or pMT3 (constitutive promoter or *clbB* promoter upstream of *luxCDABE* respectively), or the plasmid without promoter as a control. Three independent bacterial cultures grown 3.5 h at 0% or 21% oxygen were placed in a 96-well microplate and photographed with a CCD camera in a dark imaging station. Atmospheric oxygen present during the image acquisition provide the oxygen required to the reaction catalyzed by the luciferase. **B.** The luminescence (normalized to the optical density at 600 nm) was measured in a plate reader following 3.5 hours culture at specified percentages of oxygen. The mean and standard error of three independent cultures are shown, with the p values of an ANOVA and Tukey’s multiple comparison test.

**Figure 5:**
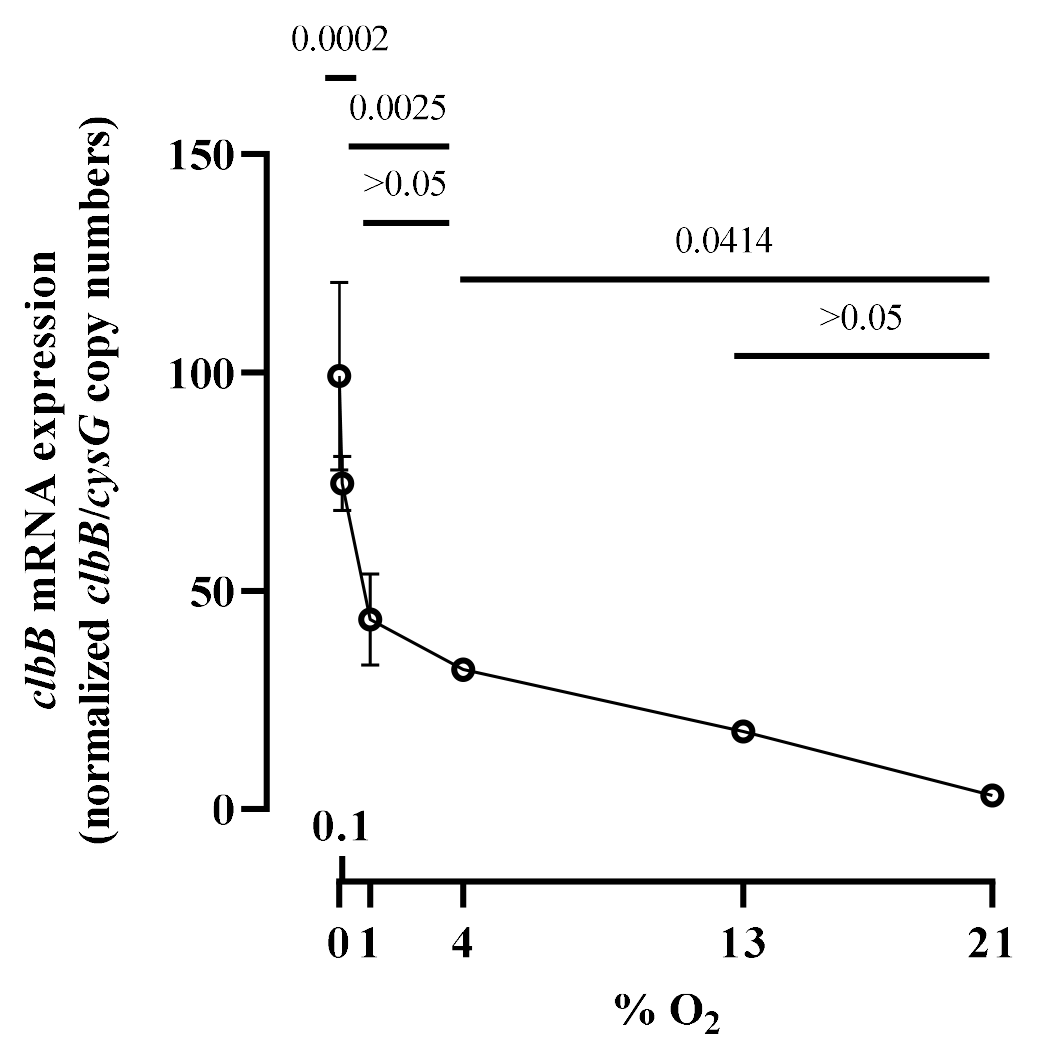
Effect of oxygen concentration on *clbB* gene expression. The mRNA levels of *clbB* gene were determined in *E. coli* SP15 grown 3.5 hours at specified percentages of oxygen. The DNA copy number of the *clbB* mRNA relative to that of the *cysG* housekeeping gene was normalized to the mean maximum level. The mean and standard error of three independent cultures are shown, with the p values of an ANOVA and Tukey’s multiple comparison test. The error bars of the biological triplicate samples at O_2_>=4% are too small to appear on the graph.

### ArcA is required for full DNA-crosslinking by *E. coli* producing colibactin

We next sought to investigate the mechanism by which oxygen regulates colibactin production. The ArcAB two component system, composed of the ArcB sensor and the response regulator ArcA, is well known to sense microaerobic and anaerobic conditions (27). To test for a role in colibactin production, we constructed a SP15 strain mutated for *arcA* and assessed its DNA cross-linking activity at a low (0.1%) oxygen concentration (Figure 6). The DNA cross-linking activity of the *arcA* mutant was significantly reduced compared to the wild-type strain, and was restored in the mutant hosting a plasmid-encoded *arcA* gene (Figure 6A and B). The growth of *E. coli* wild-type strain SP15, *arcA* mutant and complemented mutant strain was similar (Figure 6C), indicating that the altered cross-linking activity of the *arcA* mutant was not due to a growth defect and reduced bacterial numbers.

**Figure 6:**
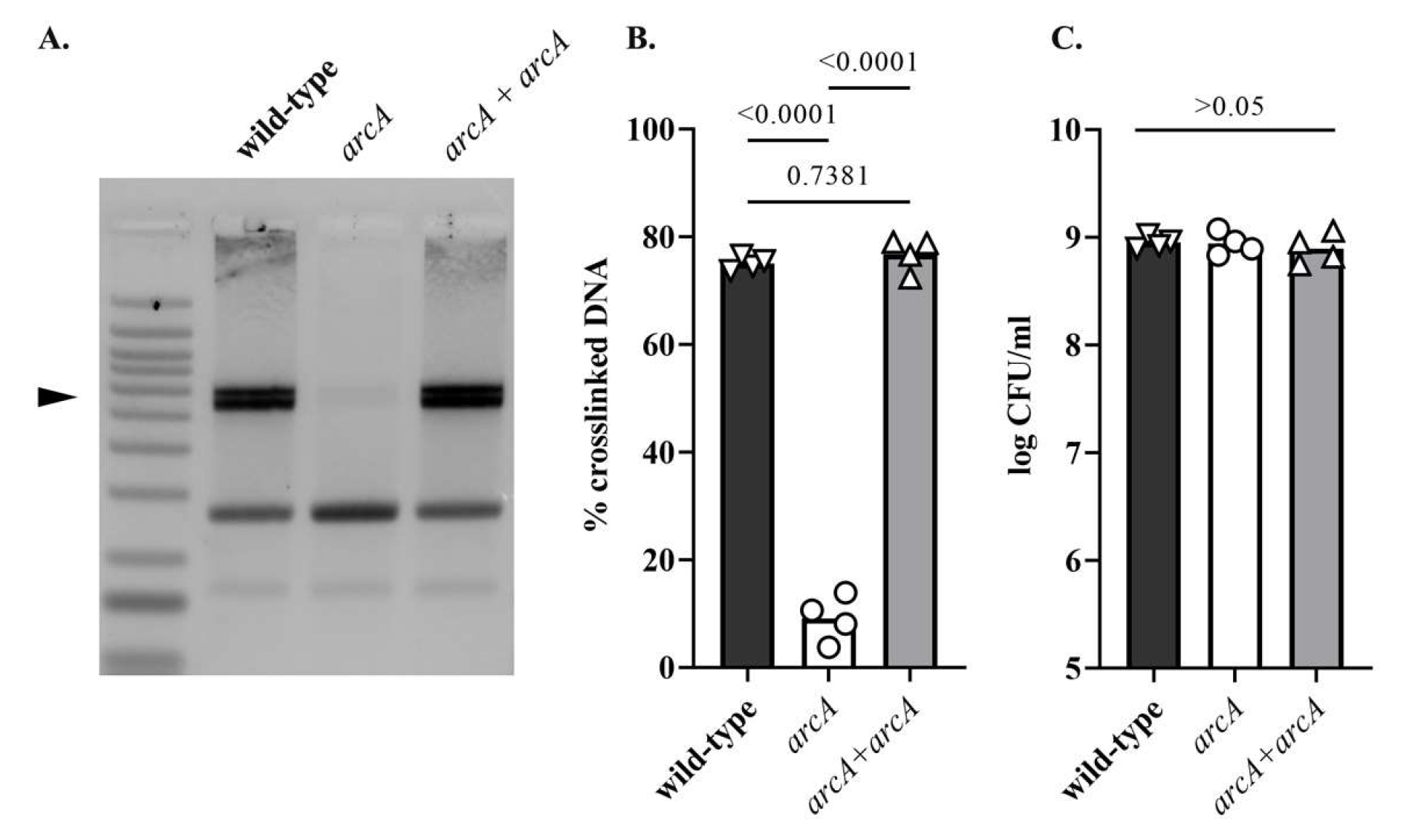
Deletion of *arcA* results in decreased DNA crosslinking activity of a *pks+ E. coli*. **A.** *E. coli* wild-type strain SP15, *arcA* mutant and *arcA* mutant complemented with a plasmid-encoded *arcA* were grown 3.5 hours at 0.1% oxygen, then incubated 40 min with linearized plasmid DNA. The DNA cross-linked by colibactin with apparent doubling in size (arrow) was visualized by denaturing gel electrophoresis. The sizes of the DNA fragments in the ladder are depicted in figure 2A. **B.** The percentage of DNA signal in the crosslinked DNA band relative to the total DNA signal in the lane was determined by image analysis. The results of four independent cultures are shown, with the p values of an ANOVA and Tukey’s multiple comparison test. **C.** In the same experiments, the bacterial growth following 3.5 hours culture was examined by plating and counting colony forming units (CFU).

### ArcA positively regulates the expression of *clbB*

The *arcA* mutant exhibited a significantly reduced *clbB* promoter activity compared to the wild-type strain (Figure 7A). In contrast, a *clbP* mutant defective in colibactin production (34) showed wild-type levels of promoter activity, indicating that colibactin production does not indirectly regulate *clbB* promoter activity. The activity of the *clbB* promoter was significantly reduced in a *clbP arcA* double mutant, as in the single *arcA* mutant (Figure 7A). However, neither the mutation of *arcA* and/or *clbP*, nor the oxygen concentration altered the activity of a constitutive control or *clbS* promoter (supplementary figure 1), indicating that ArcA specifically affects ClbB expression. We confirmed by Q-PCR that *clbB* mRNA expression was significantly reduced in the *arcA* mutant or the *clbP arcA* double mutant (Figure 7B). These results suggest that ArcA positively regulates the *clbB* gene under low oxygen conditions.

**Figure 7:**
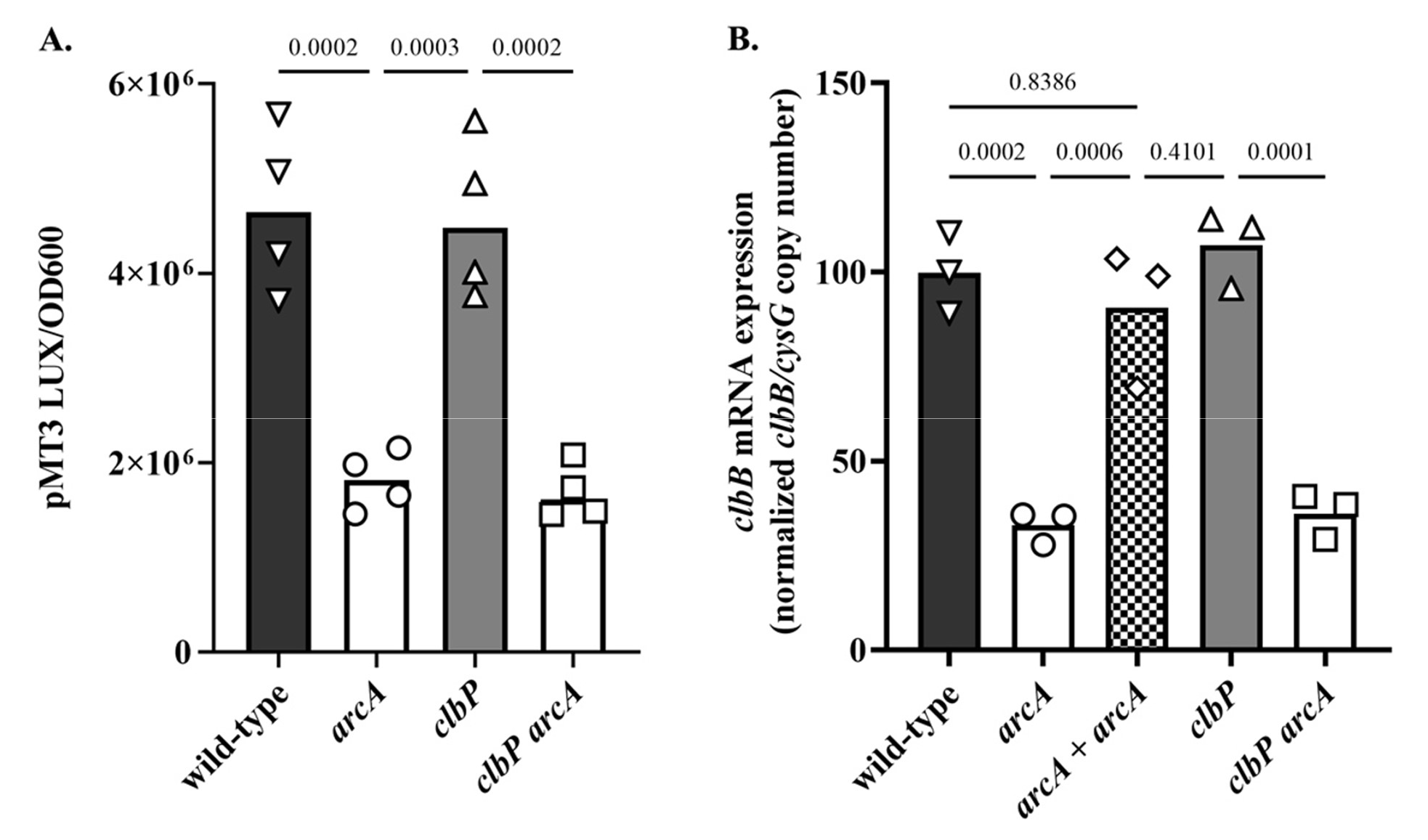
Effect of *arcA* mutation on *clbB* promoter activity and *clbB* expression. **A.** Luminescence of the *E. coli* wild-type strain SP15, the *arcA* mutant, *clbP* mutant or *clbP arcA* double mutant carrying the luciferase reporter plasmid pMT3 (encoding the promoter region of *clbB* upstream of *luxCDABE*) grown 3.5 h at 0.1% O_2_. The luminescence (normalized to the optical density at 600 nm) of four independent bacterial cultures grown 3.5 h at 0.1% O_2_ was measured in a plate reader. The mean and individual values are shown, with the p values of an ANOVA and Tukey’s multiple comparison test. **B.** The mRNA levels of the *clbB* gene were determined in *E. coli* wild-type strain SP15, the *arcA* mutant, the complemented *arcA* mutant (*arcA* + *arcA*), the *clbP* mutant or *clbP arcA* double mutant grown 3.5 h at 0.1% O_2_. The DNA copy number of the *clbB* mRNA relative to that of the *cysG* housekeeping gene was normalized to the mean wild-type level. The mean and individual values of independent cultures are shown, with the p values of an ANOVA and Tukey’s multiple comparison test.

To determine whether this regulation is direct or indirect, we searched for ArcA-binding sites within the *clbB* promoter region. Using the *in silico* PRODORIC virtual footprint tool (35), a putative ArcA binding site was located at position −39 (TGTTAAATAA, score 0.99) upstream of the ClbB open reading frame. We next performed an electrophoretic mobility shift assay (EMSA) using a DNA fragment containing the *clbB* promoter region and purified ArcA protein, which was phosphorylated or not. ArcA is converted to its active form by phosphorylation by ArcB to allow binding to the promoter region of target genes (27). Phosphorylated ArcA induced a dose-dependent band shift of the *clbB* DNA probe (Figure 8). In contrast, non-phosphorylated ArcA induced a much smaller band shift, in agreement with similar observations at other ArcA-regulated promoters (36). A similar small non-specific band shift was observed with a DNA fragment amplified from the coding region of *cysG* as a negative control, while the promoter region of *mdh* which is a known ArcA-binding locus (37), showed a strong band-shift similar to that of the *clbB* probe when bound to phosphorylated ArcA (Figure 8). Taken together, these results indicate that ArcA regulates the production of colibactin in a positive and direct manner.

**Figure 8:**
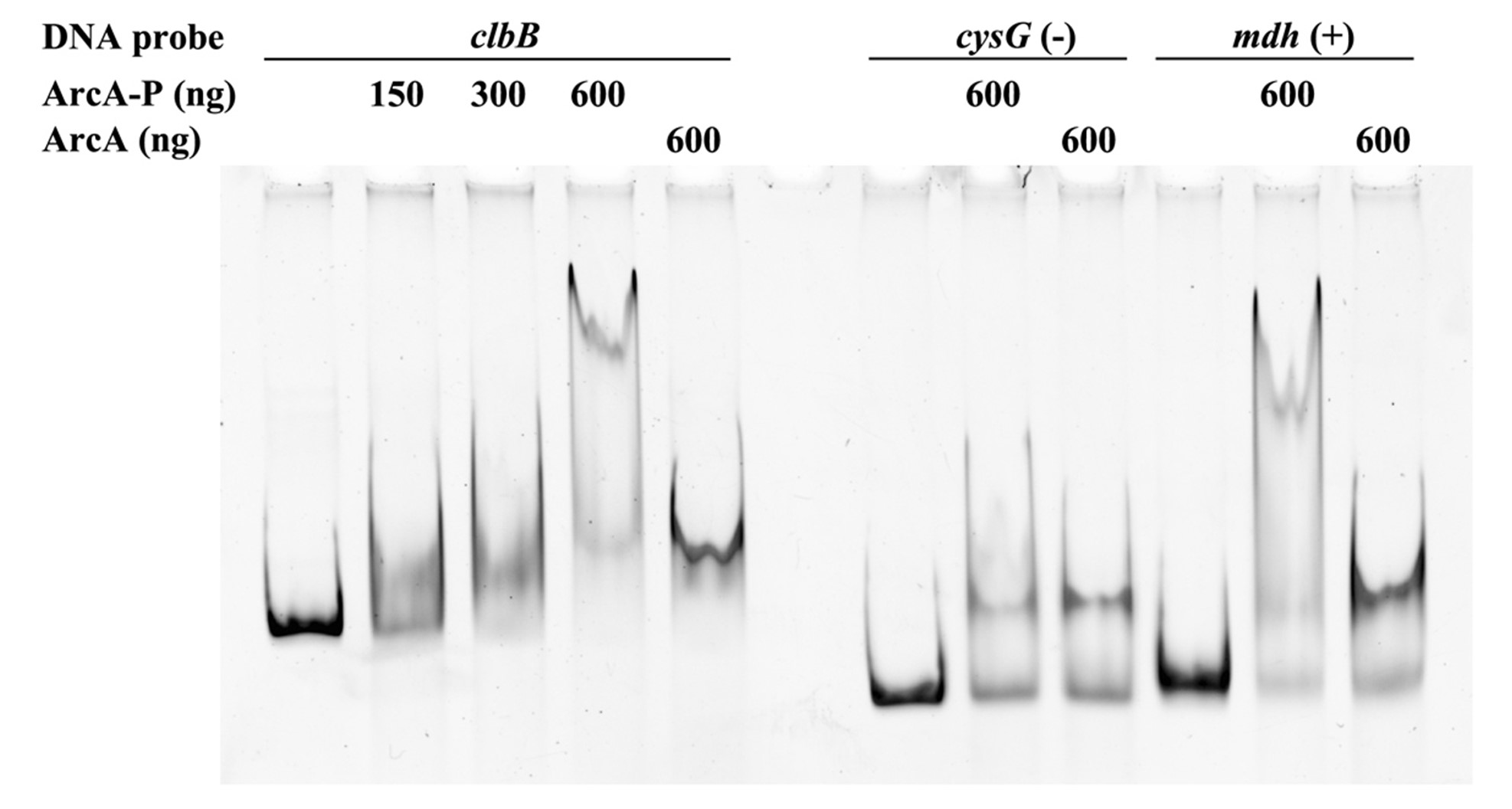
phosphorylated ArcA binds to *clbB* promoter region. EMSA using a DNA probe carrying the *clbB* promoter region and purified ArcA-P (phosphorylated *in vitro*) or ArcA, in indicated amounts. DNA probes carrying a *cysG* intragenic region or *mdh* promoter region were used as negative and positive controls, respectively.

### Oxygen and *arcA* mutation inhibit the genotoxicity of *pks+ E. coli* infecting epithelial cells

To confirm that oxygen inhibits colibactin production, we examined the genotoxicity of the *E. coli* strain SP15 in epithelial cells grown in a standard atmosphere (21% O_2_) or in “physioxic” conditions (4% O_2_) (38). Human epithelial HeLa cells were infected 4 hours with the bacteria, at 21% or 4% O_2_, then DNA damage was visualized 4 hours later by immunofluorescence staining of histone H2AX S139-phosphorylation (γH2AX) (1) (Figure 9). Infection with SP15 at 21% oxygen inflicted detectable, low-level DNA damage, while infection at 4% oxygen induced markedly higher DNA damage (Figure 9AB). HeLa cells treated with genotoxic cisplatin under 21% or 4% oxygen conditions had similar γH2AX levels, indicating that a modulation of the DNA damage response by oxygen cannot account for the increased genotoxicity of SP15 at 4% oxygen (supplementary figure 2). We also assessed the genotoxicity of the *arcA* mutant in physioxic conditions. The mutant induced significantly reduced DNA damage, while the complemented mutant had its genotoxicity restored (Figure 9AB). In contrast, the *clbP* mutant and *clbP arcA* double mutant were both non-genotoxic, with cells exposed to these mutants showing baseline levels of γH2AX (Figure 9AB).

**Figure 9:**
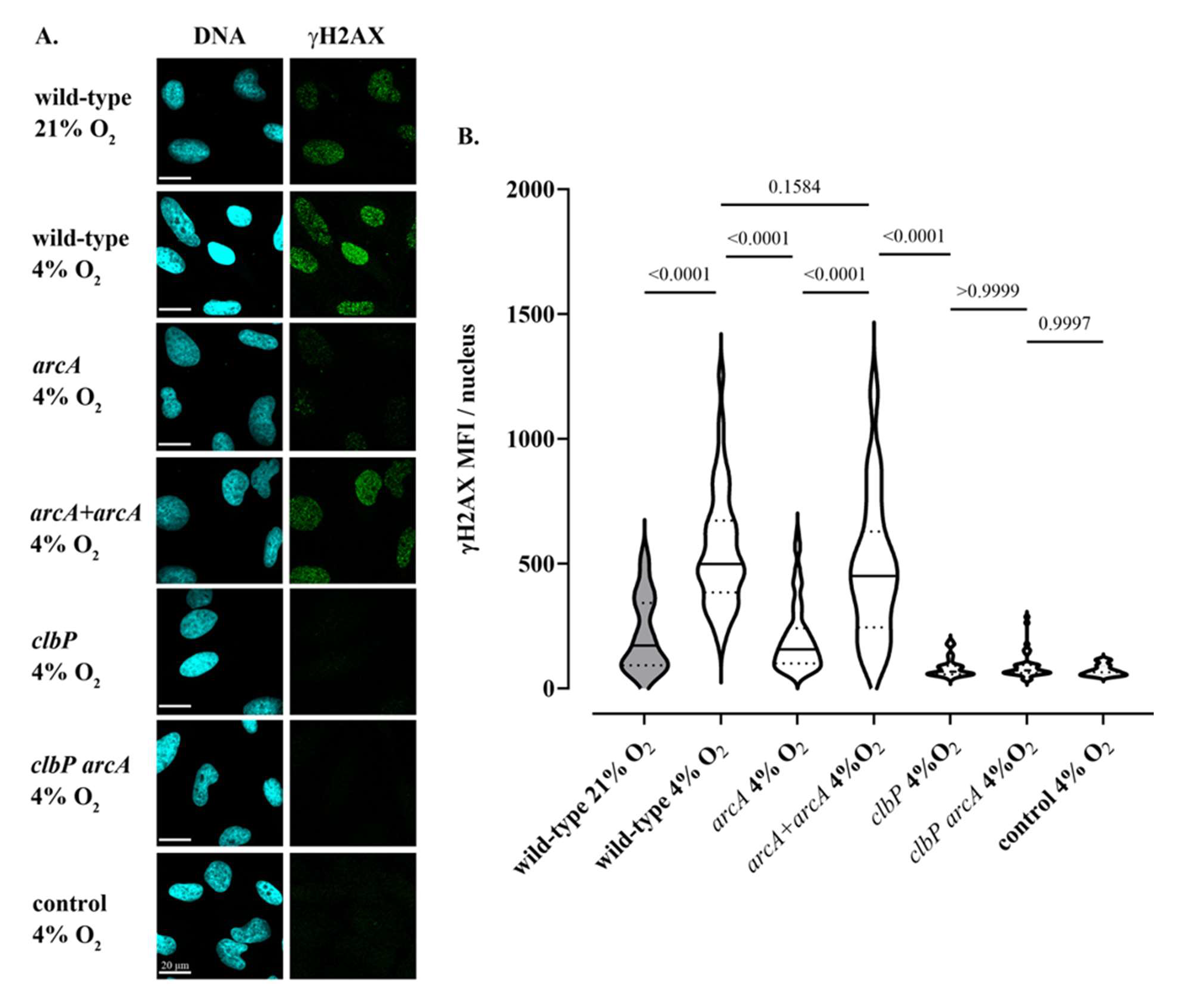
Oxygen and the *arcA* mutation inhibit the genotoxicity of colibactin-producing *E. coli*. **A.** HeLa epithelial cells were infected with *E. coli* wild-type strain SP15, the *clbP* mutant, the *arcA* mutant, the *arcA* mutant complemented with plasmid-encoded *arcA,* or the *clbP arcA* double mutant for 4 hours under 21% or 4% oxygen atmosphere, and then cellular DNA-damage was detected by γH2AX immunofluorescent staining. DNA was counterstained with DAPI. Scale bar = 20 µm **B.** γH2AX mean fluorescence intensity (MFI) within approximatively 40 to 90 cell nuclei in 2 to 4 independent experiments was measured by image analysis. The violin plots show the frequency distribution of the data, with the median and quartiles as solid and dotted lines. P values were calculated with an ANOVA and Tukey’s multiple comparison test.

To strengthen these results, the genotoxicity of the wild-type and mutant strains was tested on non-transformed rat intestinal epithelial cells (IEC-6) at 21 or 4% O_2_. The results were similar to those in HeLa cells, with an increase in the genotoxicity of *E. coli* SP15 at 4% compared to 21% O_2_, which was reduced in the *arcA* mutant, and abolished in the *clbP* and *clbP arcA* mutants (supplementary figure 3). We also examined the cytotoxicity of *pks+ E. coli* in HeLa and IEC-6 cells at 21 or 4% O_2_. Indeed, the cellular response to DNA damage induced by colibactin leads to proliferation arrest, cell death and senescence, resulting in reduced cell numbers and the formation of giant cells (megalocytosis) (13). In both cell lines, this cytotoxicity phenotype induced by the wild-type and mutant strains followed that of the genotoxicity observed with the same strains; the cytotoxicity of the wild-type SP15 was greater in physioxia, and markedly reduced in the *arcA* mutant, whereas it was restored in the complemented *arcA* mutant, but completely abolished in the *clbP* mutant or the *clbP arcA* double mutant (supplementary figure 4).

Overall, our data show that oxygen concentration modulates colibactin production and genotoxic activity of *pks*+ *E. coli* on epithelial cells via an *arcA*-dependent mechanism.

## Discussion

We show that oxygen reduces the production and genotoxicity of colibactin, and that the Arc system which sense oxygen availability positively regulates colibactin production. This regulation of colibactin production is consistent with the observation that fitness and virulence systems of enterobacteria are usually regulated by oxygen (27). Indeed, commensal as well as pathogenic enterobacteria encounter oxygen levels that are generally hypoxic, with steep gradients and dynamic oxygenation cycles. Natural oxygenation conditions are in sharp contrast to the constant, high oxygen concentration used in standard "normoxic" (21% O_2_) *in vitro* experiments. In contrast, the intestine exhibit longitudinal and cross-sectional gradients of oxygen concentration, decreasing from the small to the large intestine, and increasing from the anoxic (<0.1% O_2_) lumen to the oxygenated (∼1% O_2_) epithelium surface (39). Intestinal oxygenation further fluctuates between rest and ingestion of nutrients, when absorptive hyperemia occurs. Besides the intestinal niche, in the host epithelial tissues and fluids (such as urine), pathogenic enterobacteria encounter “physioxic” oxygen levels (2% to 9%, averaging ∼4%) (38, 40). Pathogenic enterobacteria such as *Shigella*, enterotoxigenic *E. coli*, avian and uropathogenic *E. coli* tightly regulate the expression of their virulence factors in relation to oxygen concentration (41–44). For *pks+ E. coli*, the regulation of colibactin production by oxygen via the Arc system could be viewed as a selected advantage in relation with the regulation of virulence and fitness within the host. Indeed, the metabolic cost of synthesizing a complex compound such as colibactin using eight huge NRPS and PKS enzymes is very high, implying that the selective pressure for their expression and maintenance must be strong (45). As colibactin is susceptible to aerobic oxidation (3), an energy saving strategy that inhibits its synthesis under environmental oxygen conditions leading to its inactivation may have been selected. Alternatively, the adaptation of colibactin bioactivity and production to a low-oxygen environment could be associated to the role of colibactin in the competition with the polymicrobial community in the anoxic intestinal lumen. Indeed, recent findings highlighted a direct role for colibactin in killing or inhibiting enteric bacteria (such as the anaerobe *Bacteroides fragilis*), in the induction of prophages in polymicrobial enteric anerobic cultures, and ultimately in shaping the gut microbiota (10, 11, 32). Our finding that an anoxic environment is most favorable for colibactin production is thus consistent with the recently proposed role of colibactin in interbacterial competition in gut lumen.

The notion that oxygen regulates colibactin production is also relevant to the question of whether, where and when colibactin inflicts DNA damage in the gut. It has been repeatedly reported that DNA damage was inflicted in enterocytes of rats and mice colonized or force-fed with different strains of *E. coli pks*+ (NC101, 192PP, M1/5, SP15, 11G5, the probiotic strain Nissle 1917) (15, 17, 19, 20, 46, 47). Keeping in mind that colibactin production is also regulated by a variety of environmental cues such as iron, polyamines, and carbon sources (22–24), the physiological hypoxia at the surface of the epithelium could constitute a favorable environment for colibactin production and activity. Consistent with this hypothesis, we observed in this work that *E. coli* SP15 induced marked DNA damage under physioxic conditions in two epithelial cell lines. Other low oxygen intestinal compartments could favor even more colibactin production, such as the stem cells that reside in a hypoxic niche (as low as 1% O_2_) at the bottom of the crypt (48). Signs of long-term persistent chromosomal instability have been reported in animals colonized with *pks+ E. coli* (46) and the mutational signature of colibactin has been found in intestinal crypts in healthy human subjects (49), suggesting that *pks+ E. coli* could leave their genotoxic imprint in stem cells. As far as is described in the literature, *E. coli pks+* have not yet been observed in the gut stem niche, but were shown to form biofilms together with the anaerobe *B. fragilis* on the colonic epithelium of colorectal cancer patients (50). It is well known that tumor tissue is hypoxic compared to healthy tissue (29), and thus could present another favorable environment for colibactin production. It is also interesting to note that inflammation appears to be a key factor in promoting colorectal tumorigenesis by *E. coli pks*+ in mouse models (18, 20). Severe inflammation can lead to local oxygen depletion, resulting from tissue necrosis and neutrophil respiratory burst (28). Such hypoxic areas can also be generated during infectious processes, when the host responds by coagulation to prevent bacterial dissemination, thus generating a rapid drop in local oxygen tension (51). Hypoxia at the level of the inflamed and infected tissues could therefore present environments favorable for colibactin production. Further study of the localization of *pks*+ bacteria in relation to host cell DNA damage and tissue inflammation will be necessary to better understand the impact of colibactin on infection and cancer.

In conclusion, we provide evidence that oxygen decreases colibactin synthesis by *E. coli* via an *arcA*-dependent regulatory mechanism, raising the hypothesis that colibactin production and genotoxic activity are adapted to the bacterial warfare in the anoxic gut and also to hypoxic infected, inflamed and tumor tissues.

## Materials and Methods

### Bacterial strains, mutants, plasmids and primers

The bacterial strains, plasmids and primers used in this study are listed in Table 1. Mutagenesis of *arcA* gene was performed by using the lambda red mutagenesis method, and confirmed by PCR, with the primers listed in Table 1. The mutant was trans-complemented with the pGEN-arcA plasmid. The plasmid pMT20 was constructed by PCR amplification of the 740 bp DNA sequence upstream of the *clbS* gene from the SP15 genome, and cloned into the reporter plasmid pCM17 encoding the *luxCDABE* operon. The resulting plasmid was verified by sequencing.

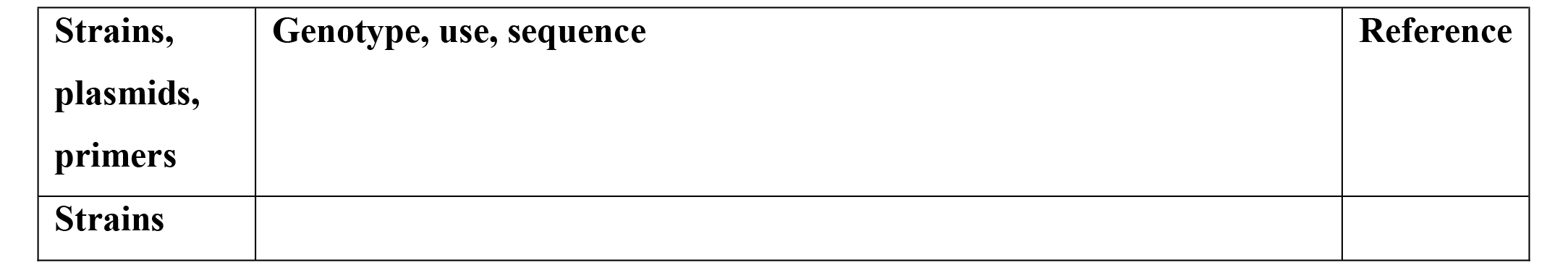
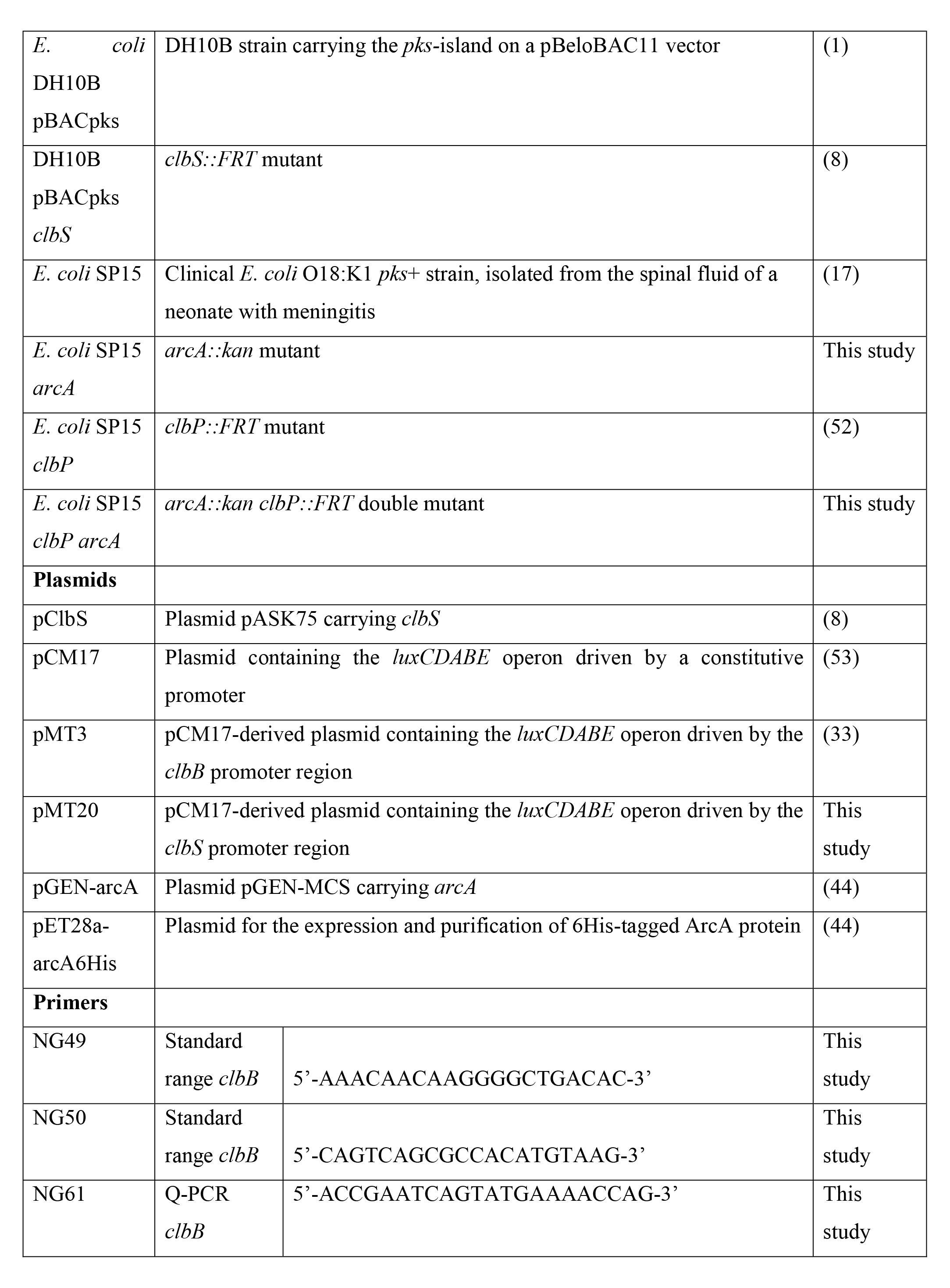
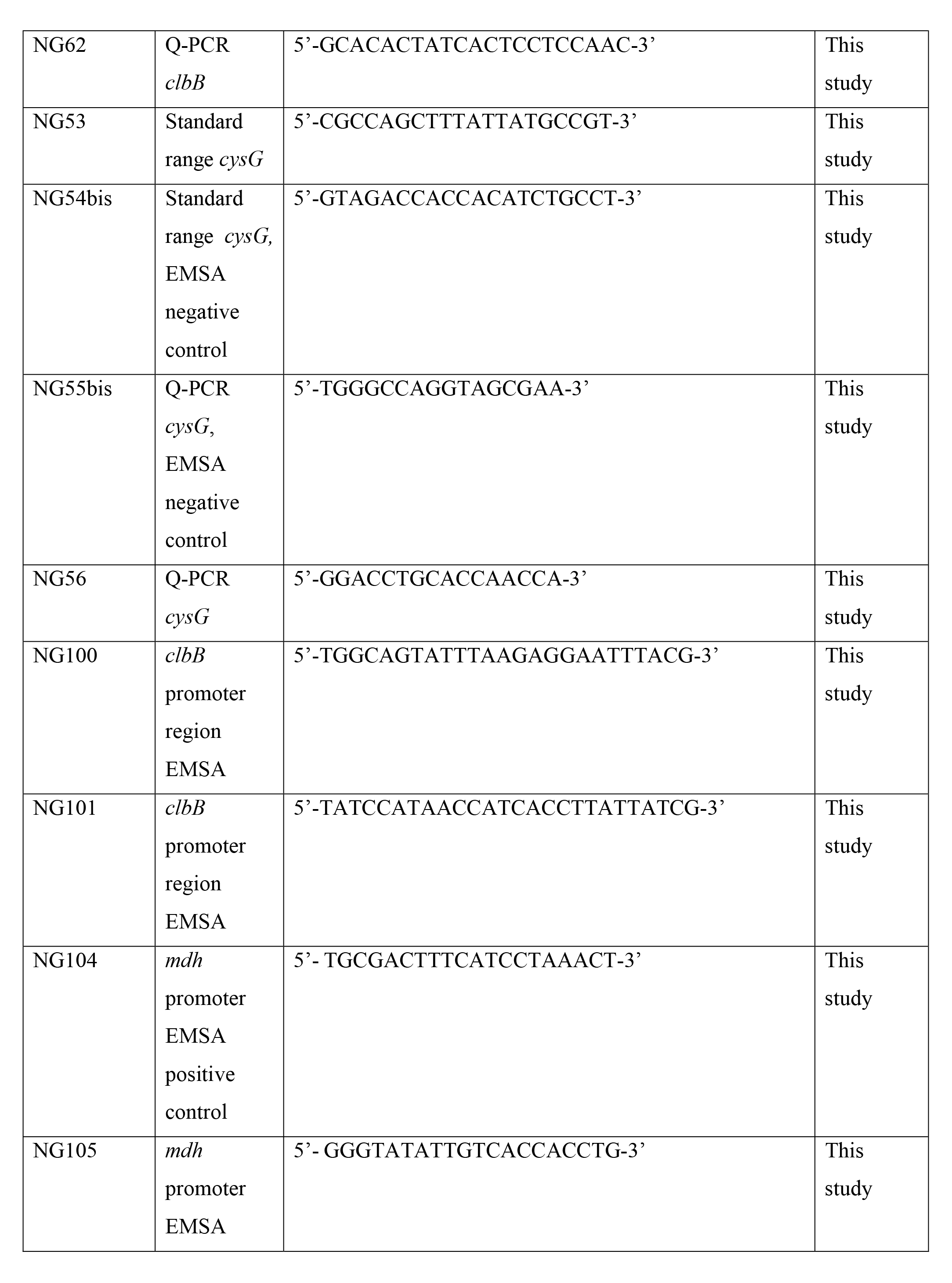
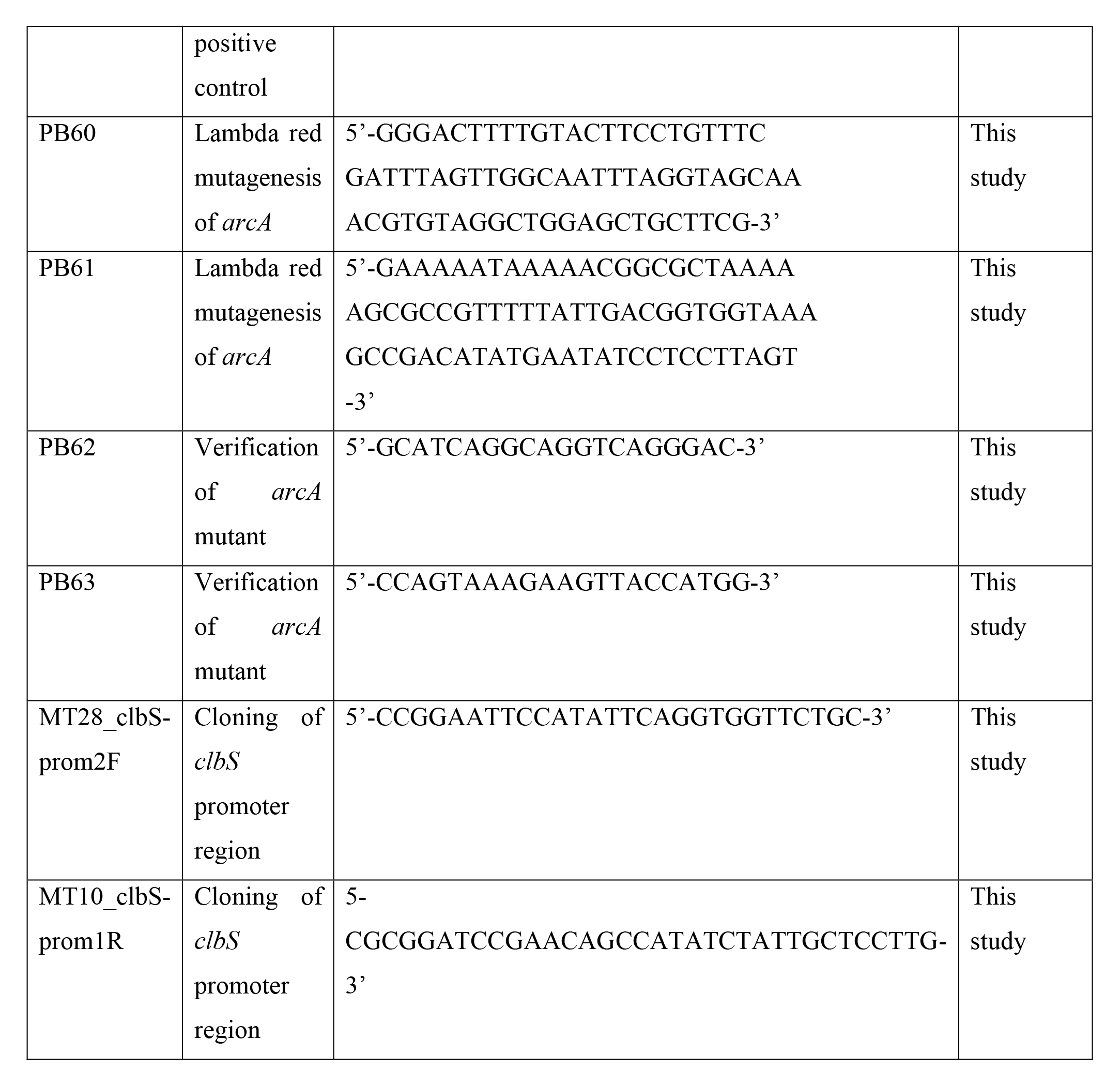

### Bacterial culture conditions

The day before the experiments, the *E. coli* strains were grown overnight in a standard incubator at 37°C with 240 RPM shaking in Lennox L broth (LB, Invitrogen) supplemented when required with appropriate antibiotics (Kanamycin 50 μg/ml, streptomycin 50 μg/ml, chloramphenicol 25 μg/ml or carbenicillin 50 μg/ml). Then, the bacteria were pre-cultured in LB or in Dulbecco’s modified Eagle’s medium (DMEM) with 25 mM HEPES (Invitrogen) to reach the exponential phase (OD600 ∼ 0.4). Cultures at various oxygen concentrations were performed in a hypoxic workstation (Whitley H35) regulated at 0.1% to 13% O_2_, 5% CO_2_, 37°C and 70-90% humidity. To achieve a 0% O_2_ anoxic atmosphere, we used a jar with anaerobic atmosphere generation bags (Sigma) manipulated within the hypoxic workstation set at 0.1% oxygen. Standard microbiological and cell culture incubators were used for the 21% O_2_ (atmospheric) condition. Culture media were equilibrated for 16-24 h at the required oxygen concentration before use. For DH10B strains (Figure 1), LB cultures were inoculated with 2×10^6^ bacteria/ml in 5 ml and grown overnight (17 h) in fully closed 10 ml tubes (Kima), or in opened 50 ml tubes (Falcon), or in 14 ml 17×100 mm tubes with the cap in vented position (Falcon 352059). The *E. coli* strain SP15 was inoculated (1.5×10^7^ bacteria) in 1 ml DMEM Hepes in Falcon 352059 tubes with the cap in vented position and cultivated 3.5 h with 240 RPM agitation at 37°C within the hypoxystation set at the oxygen concentration required. For CFU enumeration, the cultures were serial-diluted in PBS, plated on LB agar plates and incubated overnight at 37°C in a standard incubator.

### Exogenous DNA cross-linking by colibactin-producing *E. coli*

The assay was performed as described before (4). Briefly, 100µl of bacterial culture were added with 500 ng DNA (pUC19 linearized by BamHI digestion) and 1mM EDTA, then incubated 40 min at the indicated oxygen concentration. The DNA was recovered using a Qiagen PCR purification kit, analyzed by denaturing gel electrophoresis, stained with GelRed (Biotium) and visualized with a Bio-Rad Chemidoc XRS system. DNA bands were quantified from unsaturated flat-fielded images using ImageJ.

### Quantification of N-myristoyl-D-AsnOH (C14-Asn)

Bacterial supernatants (1.5 ml) were prepared by centrifugation at 5000 g, filtration through 0.22 µm PVDF filters (Millipore) and stored at −80°C. Lipids were extracted and analyzed as before (54): briefly, 1 ml of sample was extracted on solid phase 96 wells plates (OASIS HLB, Waters), eluted with methanol, evaporated twice under N_2_ then suspended in 10 µl methanol. The quantification of C14-Asn was performed by the MetaToul Lipidomics facility (Inserm UMR1048, Toulouse, France), using an in-house quantification assay by high-performance liquid chromatography/tandem mass spectrometry analysis.

### Bioluminescence monitoring of *clbB* and *clbS* promoter activity

The *E. coli* strain SP15 hosting pMT3, pMT20 or pCM17 (containing the *Photorhabdus luminescens luxCDABE* operon driven by the *clbB* or *clbS* promoter region (33) or a constitutive promoter (53) respectively) was cultivated at the required concentration of oxygen, then 100µl were transferred on 96 wells black plates (Greiner), the plate was shaken vigorously 5 sec to provide oxygen to the luciferase and the bioluminescence was measured with a Tecan Spark plate reader or a Perkin Elmer EnSight multimode plate reader, or photographed with a Bio-Rad Chemidoc XRS system.

### *clbB* mRNA quantification

10^9^ bacteria were collected by centrifugation and stored at −80°C. RNA was purified using the RNA Easy plus mini kit (Qiagen), then 2 µg were used to synthesize cDNA with the i-script kit (Bio-Rad). For standard range of *clbB* and *cysG* housekeeping gene (55), PCRs were performed using *E. coli* SP15 genome as a template and the primer pairs NG49/NG50 and NG53/NG54 respectively. Standards were purified and serial diluted from 5.10^9^ to 5.10^1^ copy/µl. Real time PCR were performed with the iQ SYBR Green supermix (Bio-Rad), with 2 µl of cDNA or standard dilutions, and 187.5 nM of primer pairs NG61/NG62 (*clbB*) or NG55/NG56 (*cysG*). The copy numbers of *clbB* and *cysG* were calculated from the Ct obtained using the standard curves. The expression of *clbB* gene at different oxygen concentration was calculated by the ratio *clbB* / *cysG* copy numbers and normalized at 100% by the mean at 0.1% oxygen.

### Epithelial cells infection, quantification of DNA-damage and megalocytosis assay

The DNA-damage and megalocytosis assays were performed as described before (4, 52) with some modifications. HeLa and IEC-6 cells were cultivated in a standard atmospheric 37°C 5% CO_2_ incubator, in DMEM Glutamax (Invitrogen) supplemented with 10% fetal calf serum (FCS), 50 µg/ml gentamicin and 1% non-essential amino acids (Invitrogen). IEC-6 cells were supplemented with 0.1 U/mL of bovine insulin (Sigma-Aldrich). The cells were seeded in 8- chambers slides or in 96-wells plates (Falcon) and grown 24 hours in the atmospheric 5% CO_2_ incubator, or in the hypoxic workstation set at 4% O_2_ 5% CO_2_. The cells were infected in DMEM 25 mM HEPES at a multiplicity of 100 bacteria per cell and incubated 4 hours in the hypoxic or standard incubator. As controls, the cells were left uninfected or were treated with 20 µM cisplatin (Sigma) to induce DNA-damage. The cells were then washed 3 times with HBSS supplemented with 100 µg/ml gentamicin, incubated in complete cell culture medium supplemented with 100 µg/ml gentamicin for 4 hours, in the same hypoxic or standard incubator. The cells were finally washed in PBS and fixed 15 min in PBS 4% formaldehyde. Following 5 min permeabilization in 0.2% Triton X-100, and blocking in MAXblock medium (Active Motif), the cells were stained with antibodies against γH2AX (1:500, JBW301, Millipore) followed with anti-mouse AlexaFluor 488 (1:1000, Invitrogen). The cells were mounted in Fluoroshield w/DAPI medium (Sigma), and examined with a Zeiss 710 laser scanning confocal microscope at 63X. The mean fluorescence intensities of γH2AX within the nuclei were analyzed using a NIH ImageJ macro: the nuclei were identified in the DNA image and the mean fluorescence intensity of γH2AX was measured in the masked green channel for each nucleus. The megalocytosis phenotype induced by colibactin was examined as previously described (52). Briefly, HeLa and IEC-6 cells in were infected as above in the standard atmospheric or hypoxic (4% oxygen) incubator, then washed and incubated with gentamicin for 48 h before staining with methylene blue. The cells were photographed using an Olympus BX41 microscope equipped with a 10x objective and a Leica DCF300FX camera.

### Purification, phosphorylation of ArcA-6His protein and electrophoretic mobility shift assay (EMSA)

ArcA-6His purification and phosphorylation were performed as described previously, with some modifications (44). Briefly, *E. coli* BL21 pET28a-arcA6His was grown overnight at 28°C in 100 ml of LB with 0.1 mM IPTG. Bacteria were lysed in equilibration buffer (20 mM NaH_2_PO_4_, 300 mM NaCl, 10 mM imidazole, pH 8) supplemented with lysozyme and anti-protease cocktail (Roche), followed by sonication. The lysate was cleared by centrifugation (10000 g, 30 min) and the supernatant was loaded 1 h on HisPur Ni-NTA superflow agarose (Fisher). The agarose was washed with equilibration buffer and eluted with increasing imidazole concentration (25, 50, 100 mM). The eluate was dialyzed in 20 mM Tris 50 mM NaCl 40 mM EDTA 4 mM dithiothreitol [DTT] 10% glycerol pH 7.4 buffer 16 h at 4°C with three buffer changes. Then the protein sample was concentrated using an Amicon Ultra centrifugal tube (Merck) with a buffer change to phosphorylation buffer 2X (100 mM Tris pH 8, 1mM EDTA, 20 mM glycerol, 20 mM MgCl_2_). The protein sample was stored at 4°C. Phosphorylation of 20 µg ArcA-6His was performed with 100 mM lithium potassium acetyl phosphate in 1X phosphorylation buffer. A non-phosphorylated protein control was prepared by omitting the lithium reagent. After incubation for 60 min at 25°C the samples were used immediately for DNA binding assay.

EMSA was performed as previously described (44). Briefly, DNA probes with the promoter regions of *mdh* as positive control, an intragenic region of *cysG* as negative control, and the promoter region of *clbB* were generated by PCR using primers listed in table 1. DNA probes were gel-purified (Qiagen kit), then 50 ng of DNA was mixed with increasing amounts of phosphorylated ArcA-His (0 to 600 ng) or non-phosphorylated protein (600 ng) in binding buffer (100mM Tris pH7.4, 10 mM MgCl_2_, 5% glycerol, 400 µg/mg BSA, 2 mM DTT, 100 mM KCl) for 30 min at 37°C. The reaction mixtures were then subjected to electrophoresis on a 6% polyacrylamide Novex TBE gel (Thermo) in 0.5× TBE Novex running buffer, at 200 V for 45 min. The gel was stained in 0.5× TBE containing 1× SYBR Gold nucleic acid stain (Life Technologies) for 30 min, and then the image was recorded using a ChemiDoc (BioRad).

### Statistical analyses

For quantifications, three or four independent experiments were performed for each oxygen concentration tested. CFU counts were log-transformed. Statistical analyses were performed with GraphPad Prism v9. P values were calculated using ANOVA followed by Tukey post-tests, or with a t-test.

## Acknowledgement

We thank Dr. Ganwu Li for the gift of the pGEN-arcA and pET28a-arcA plasmids, and Simon Lachambre & Lhorane Lobjois for technical assistance at the Infinity Cell Imaging Facility, Toulouse.

## Data availability statement

The authors confirm that the data supporting the findings of this study are available within the article.

---

**Supplementary figure 1:**
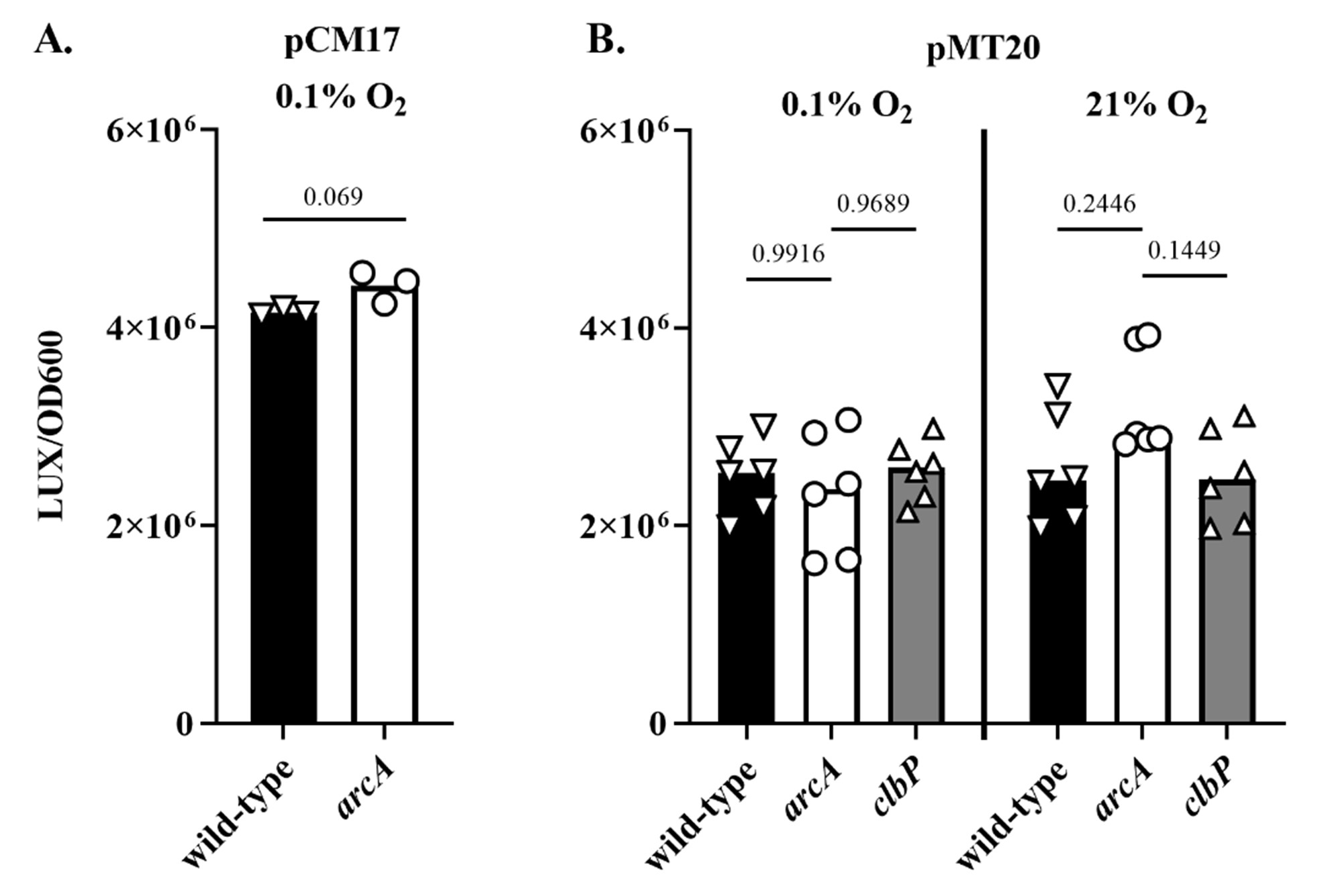
**A.** Luminescence of *E. coli* strain SP15 carrying the luciferase reporter plasmid pCM17 (constitutive promoter upstream of *luxCDABE*) grown 3.5 h at 0.1% oxygen. The mean and individual values of three independent cultures are shown, with the p value of a t-test. **B.** Luminescence of *E. coli* strain SP15 carrying the luciferase reporter plasmid pMT20 (promoter region of *clbS* upstream of *luxCDABE*) grown 3.5 h at 0.1% or 21% oxygen. The mean of 6 independent cultures are shown, with the p values of an ANOVA and Tukey’s multiple comparison test.

**Supplementary figure 2:**
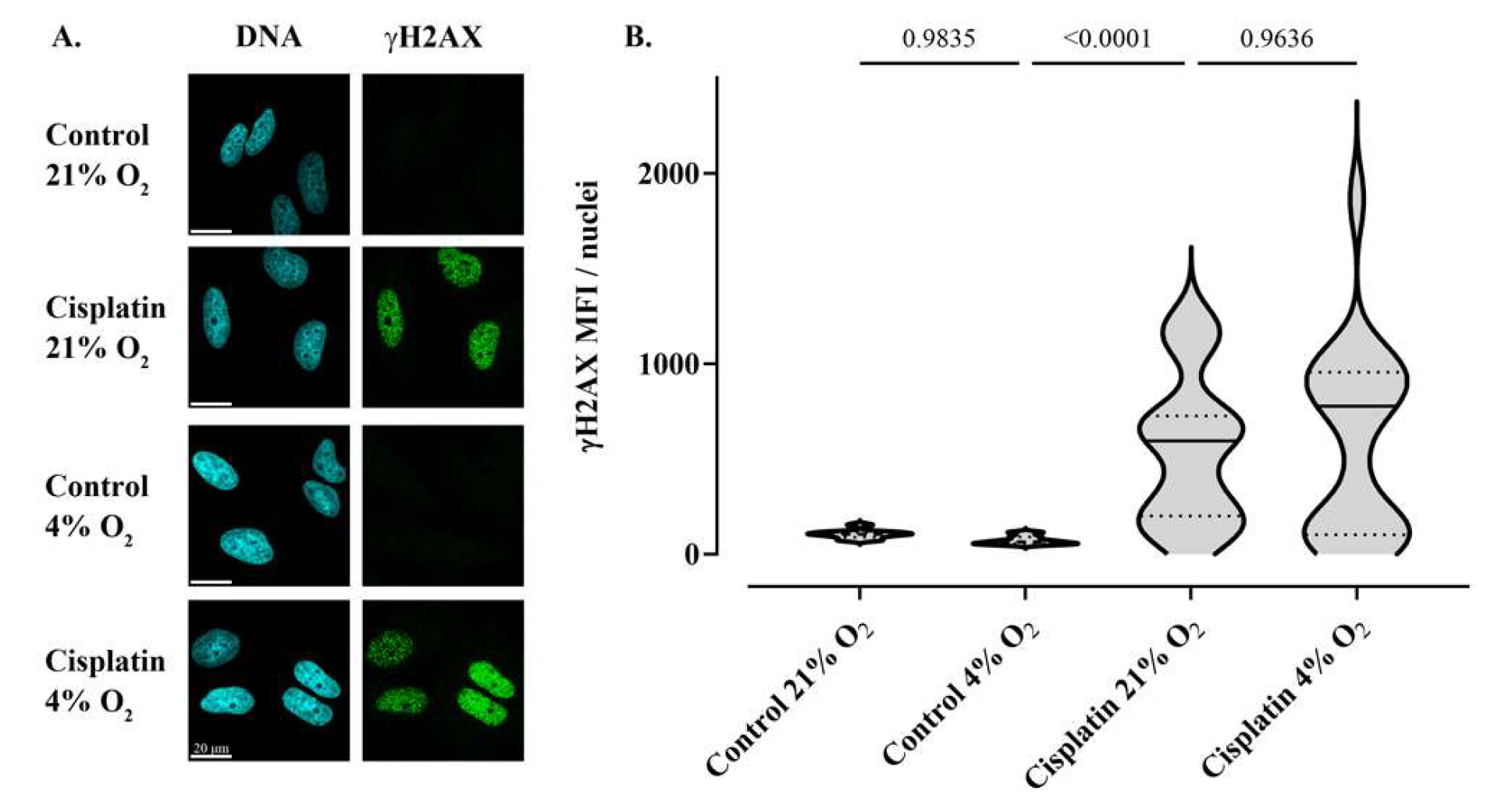
Oxygen does not alter the γH2AX response of genotoxin-treated HeLa cells. **A.** HeLa epithelial cells were treated with cisplatin for 4 hours under 21% (“hyperoxic”) or 4% (“normoxic”) oxygen atmosphere, and then cellular DNA-damage was demonstrated by γH2AX immunofluorescent staining. DNA was counterstained with DAPI. Scale bar = 20 µm **B.** γH2AX mean fluorescent intensity (MFI) within ∼50 cell nuclei was measured by image analysis. The violin plots show the frequency distribution of the data, with the median and quartiles as solid and dotted lines. P values were calculated with an ANOVA and Tukey’s multiple comparison test.

**Supplementary figure 3:**
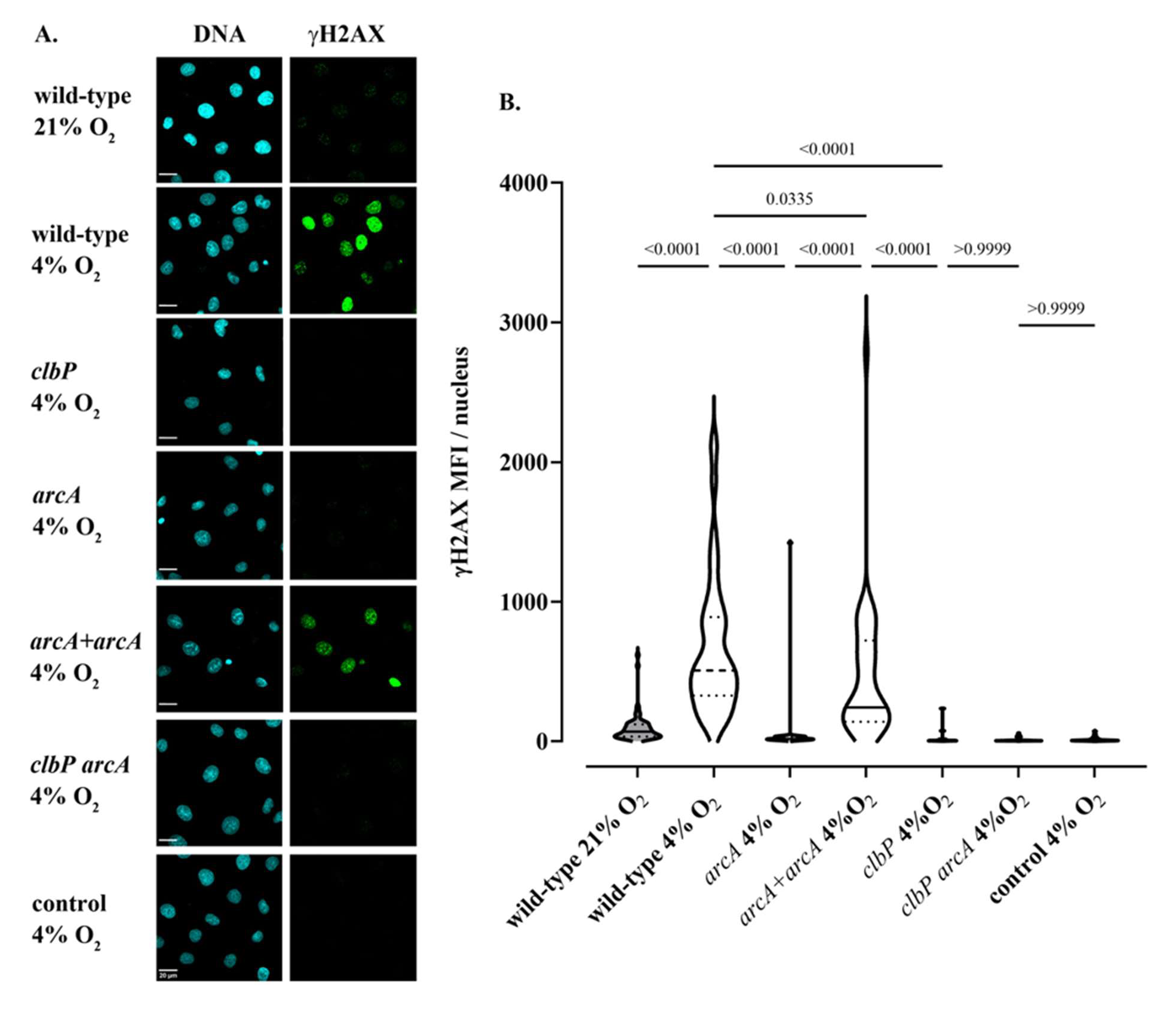
High oxygen concentration and *arcA* deletion inhibit the genotoxicity of colibactin-producing *E. coli* in intestinal IEC-6 cells. **A.** Non-transformed rat intestinal epithelial IEC-6 cells were infected with *E. coli* wild-type strain SP15, the *clbP* mutant, the *arcA* mutant, the *arcA* mutant complemented with plasmid-encoded *arcA,* or the *clbP arcA* double mutant for 4 hours under 21% or 4% O_2_ atmosphere and then cellular DNA-damage was detected by γH2AX immunofluorescence staining. DNA was counterstained with DAPI. Scale bar = 20 µm **B.** γH2AX mean fluorescence intensity (MFI) within approximatively 60 nuclei was measured by image analysis. The violin plots show the frequency distribution of the data, with the median and quartiles as solid and dotted lines. P values were calculated with an ANOVA and Tukey’s multiple comparison test.

**Supplementary figure 4:**
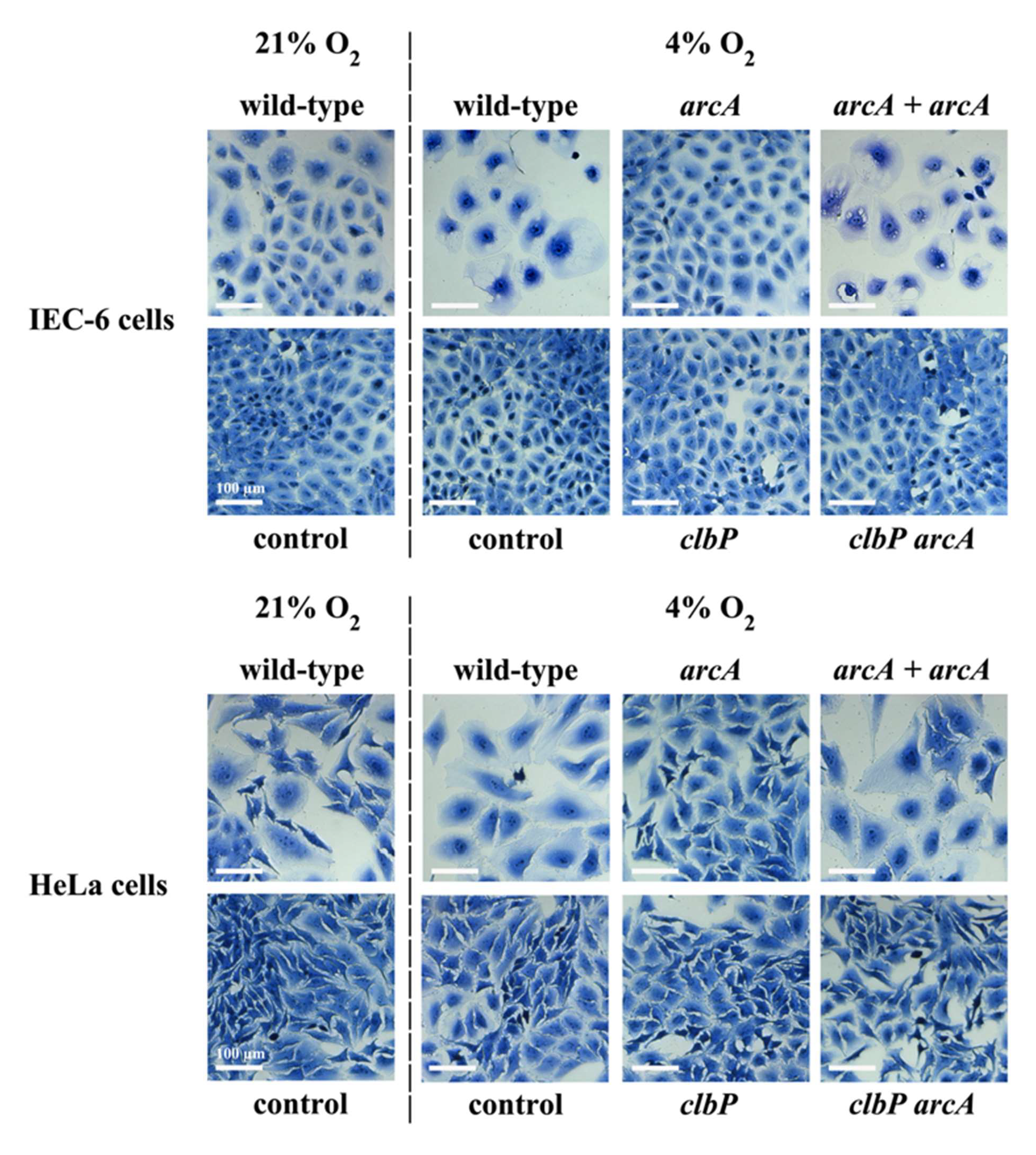
Cytotoxicity phenotype observed in IEC-6 and HeLa cells 48 h after infection with *E. coli* wild-type strain SP15, the *arcA* mutant, the *arcA* mutant complemented with plasmid-encoded *arcA,* the *clbP* mutant or the *clbP arcA* double mutant, under 21% or 4% O_2_ atmosphere. Control cells were left non-infected. Scale bars = 100 µm.

